# Image-based Cell Phenotyping Using Deep Learning

**DOI:** 10.1101/817544

**Authors:** Samuel Berryman, Kerryn Matthews, Jeong Hyun Lee, Simon P. Duffy, Hongshen Ma

## Abstract

The ability to phenotype cells is fundamentally important in biological research and medicine. Current methods rely primarily on fluorescence labeling of specific markers. However, there are many situations where this approach is unavailable or undesirable. Machine learning has been used for image cytometry but has been limited by cell agglomeration and it is unclear if this approach can reliably phenotype cells indistinguishable to the human eye. Here, we show disaggregated single cells can be phenotyped with a high degree of accuracy using low-resolution bright-field and non-specific fluorescence images of the nucleus, cytoplasm, and cytoskeleton. Specifically, we trained a convolutional neural network using automatically segmented images of cells from eight standard cancer cell-lines. These cells could be identified with an average classification accuracy of 94.6%, tested using separately acquired images. Our results demonstrate the potential to develop an “electronic eye” to phenotype cells directly from microscopy images indistinguishable to the human eye.

## Introduction

Cancer cell lines have been extensively used to model the disease as well as to screen for potential therapeutic agents. However, since cancer is such a heterogeneous condition, the ultimate utility of these cancer cell line models depends on the ability to accurately classify them. Cancer cell lines are primarily classified based on the histopathology of the original tumor but subtyping of the cell lines may require lengthy and significant molecular and genetic profiling^1–4^. Immunofluorescence phenotyping has contributed significantly to reducing the burden for cell line classification but immunofluorescence relies on the expression of specific cell surface antigens, which is expensive and error-prone, despite significant efforts in standardizing staining, data collection and automation of analysis^5^. Specifically, immunofluorescence phenotyping may be undesirable because: (1) phenotyping markers may be unavailable or lack specificity, (2) the sample may be too heterogeneous, (3) number of markers required may exceed the number of available fluorescence channels that can be detected, and (4) specific labeling may affect the cell in undesirable ways, such as activation or loss of viability. In many of these situations, an important question is whether individual cells could be phenotyped directly using microscopy images without specific labeling.

Previous approaches for image-based cell phenotyping typically rely on manual feature engineering, which involves extracting specific image features from each cell, such as size, shape, and texture of the nucleus and cytoplasm^6,7^. Machine learning approaches, such as support vector machines, are then used to classify cells using these features. The ability to identify specific phenotypes using this approach is limited because the features are defined arbitrarily and not necessarily biologically relevant. Additionally, feature extraction from microscopy images is a highly variable process that depends on many manually tuned parameters. In order to address these issues, machine learning approaches have been used to phenotype cells directly using microscopy images^8,9^. However, previous phenotyping studies have been restricted to broad cell groups, such as lymphocytes, granulocytes, and erythrocytes, that have morphologies easily distinguishable to the human eye^10–13^. Phenotyping cells with more subtle morphologies has been largely restricted to binary classification in order to detect specific alterations resulting from disease^10,14–17^. Another approach is to use brightfield microscopy images to predict the location of immune-stains on sub-cellular structures in order to identify organelles^18–21^. However, a further step is required to interpret these stains to establish the cell phenotype.

A key challenge in developing machine learning algorithms for classifying cells from microscopy images is segmenting larger microscopy fields into single-cell images. Specifically, adhesion cells are notoriously difficult to segment because they grow next to one another making their boundaries difficult to distinguish^22–24^. This segmentation problem could be dramatically simplified by enzymatically disaggregating cells prior to imaging. However, it is currently unclear if phenotypic information is sufficiently preserved in disaggregated cells.

Here, we show disaggregated cells can be phenotyped with a high degree of accuracy from bright-field and non-specifically stained microscopy images using a trained convolutional neural network (CNN). We trained the CNN using 75×75 pixel brightfield and fluorescence microscopy images of individual cells from eight standard cancer cell lines. All cells were non-specifically stained to identify their nucleus (Hoechst), cytoplasm (Calcein), and cytoskeleton (SiR-actin). The resulting images were non-distinguishable by the human-eye, however, the CNN achieved a cross-validation accuracy of 96.0±0.8%. We further tested the network using separately prepared cell images and achieved an average class accuracy of 94.6%. This work demonstrates the potential to develop an “electronic eye” to phenotype both adherent and suspension cells interchangeably, based on brightfield and non-specific fluorescence images without the need for specific markers.

## Results

### Imaging

To acquire a training set for deep-learning, we imaged 8 standard cancer cell lines. These cell lines were derived from colorectal (HCT-116) and prostate (PC3) carcinoma, prostate (LNCaP) and mammary (MCF7) adenocarcinoma, osteosarcoma (U2OS), as well as neutrophilic (HL60), monocytic (THP-1) and T-cell (Jurkat) leukemia. All cells were seeded in 96-well imaging plates and stained using DAPI to visualize the nucleus, Calcein-AM to assess cell viability and cytoplasmic morphology via intracellular esterase activity and SiR-Actin incorporation to visualize cytoskeletal morphology. Following staining, the cells were imaged using a 10X objective on a Nikon Ti-2E microscope and DS-Qi2 camera, to acquire fields of 2424×2424 pixels.

### Segmentation

We developed a Python program to extract 75×75 pixel images of individual cells from wide-field microscopy images. Our program first identified the locations of individual cells by identifying cell nuclei using an Otsu threshold on the DAPI channel (**Fig. 1a**). Small 75×75 pixel images centered on each of the nuclei were then cropped out and fed through a series of tests to reject problematic images. Firstly, cells that did not stain with Calcein-AM were rejected as non-viable. Second, cells that presented with multiple DAPI-positive nuclei or overlapping cells, debris or other imaging artefacts were also rejected. Finally, a minimum fluorescence emission threshold was used to ensure adequate signal intensity for image analysis. Together, this digital processing produced a set of high-quality images for each cell with strong staining for each of the three dyes (**Fig. 1b**). The total number of successfully segmented images for each cell type is listed in **Table 1**.

**Table 1.** Size of each class in the training and testing sets.

| CLASS | TRAINING | TESTING |
| --- | --- | --- |
| HCT-116 | **7151** | **3306** |
| HL60 | **26298** | **1313** |
| JURKAT | **9208** | **3980** |
| LNCAP | **2555** | **1216** |
| MCF7 | **4109** | **1702** |
| PC3 | **4700** | **1908** |
| THP-1 | **8656** | **1371** |
| U2OS | **15413** | **1069** |

**Figure 2.**
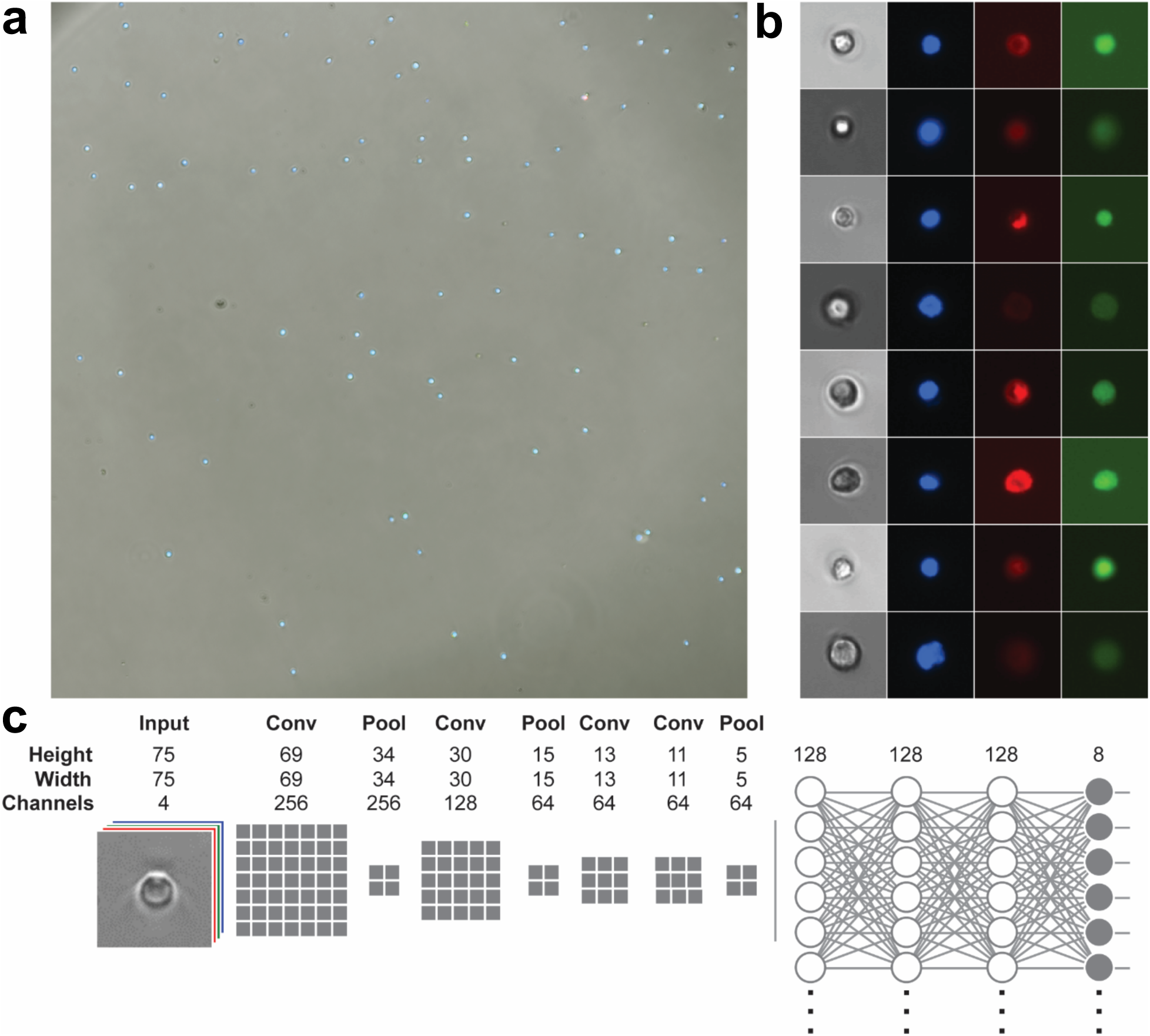
Segmentation and network design. **a,** Sample widefield microscopy image taken on a Nikon TI-2E with a QI2 Camera. The three fluorescent channels are showing nuclear (Hoechst), cytoskeleton (SiR-Actin) and cytoplasm (Calcein). **B,** Sample segmented 75×75 single cell images. Channels shown from left to right are brightfield, Hoechst, SiR-Actin and Calcein. **c,** Network model where the number of grey boxes represents the size of the kernel being used for each convolution layer with a stride of one. Not shown in the image is that after every convolution or fully-connected layer is a ReLU activation and batch normalization as well as dropout of 20% after each fully-connected layer.

### Network design

We designed a CNN consisting of a feature extraction and a classification section using the Keras library in TensorFlow. The feature extraction section consists a series of 4 convolution layers with 3 max-pooling layers. We investigated a range of initial layer kernel sizes, as a larger kernel size is expected to capture more robust features. However, we ultimately selected a kernel size of 7×7 based on the observation that larger kernel sizes did not significantly improve performance. Each convolution layer was followed by batch normalization and ReLU activation. The classification section of the network consisted of 3 fully connected layers followed by a smaller fully connected output layer. Each of the 3 fully connected layers was followed by batch normalization, ReLU activation and 20% dropout. The output of the model used a SoftMax error function for backpropagation during training.

### Training

The training accuracy of a network typically sets a ceiling on the expected outcome of validation accuracy. Therefore, the goal of network design is to reduce the gap between training and validation accuracy. We trained our CNN using a balanced dataset of 10,000 images from each cell phenotype. The dataset was balanced to avoid biasing training results. Classes containing >10,000 cell images were sub-sampled, while classes containing < 10,000 cell images were augmented using random integer multiplications of 90-degree rotations. The network was trained over 25 epochs on the dataset. The network was able to train quickly with no volatility due to pairing batch normalization with a small amount of dropout between layers (**Fig. 2a**). The batch normalization prevented runaway weights and reduced model training time while dropout prevented convergence on local minimums. The final training accuracy for the 4-channel model was 98.2%, while the training accuracies were lower for individual channels of brightfield (92.2%), nucleus (91.6%), cytoplasm (95.0%), and cytoskeleton (95.3%).

**Figure 2.**
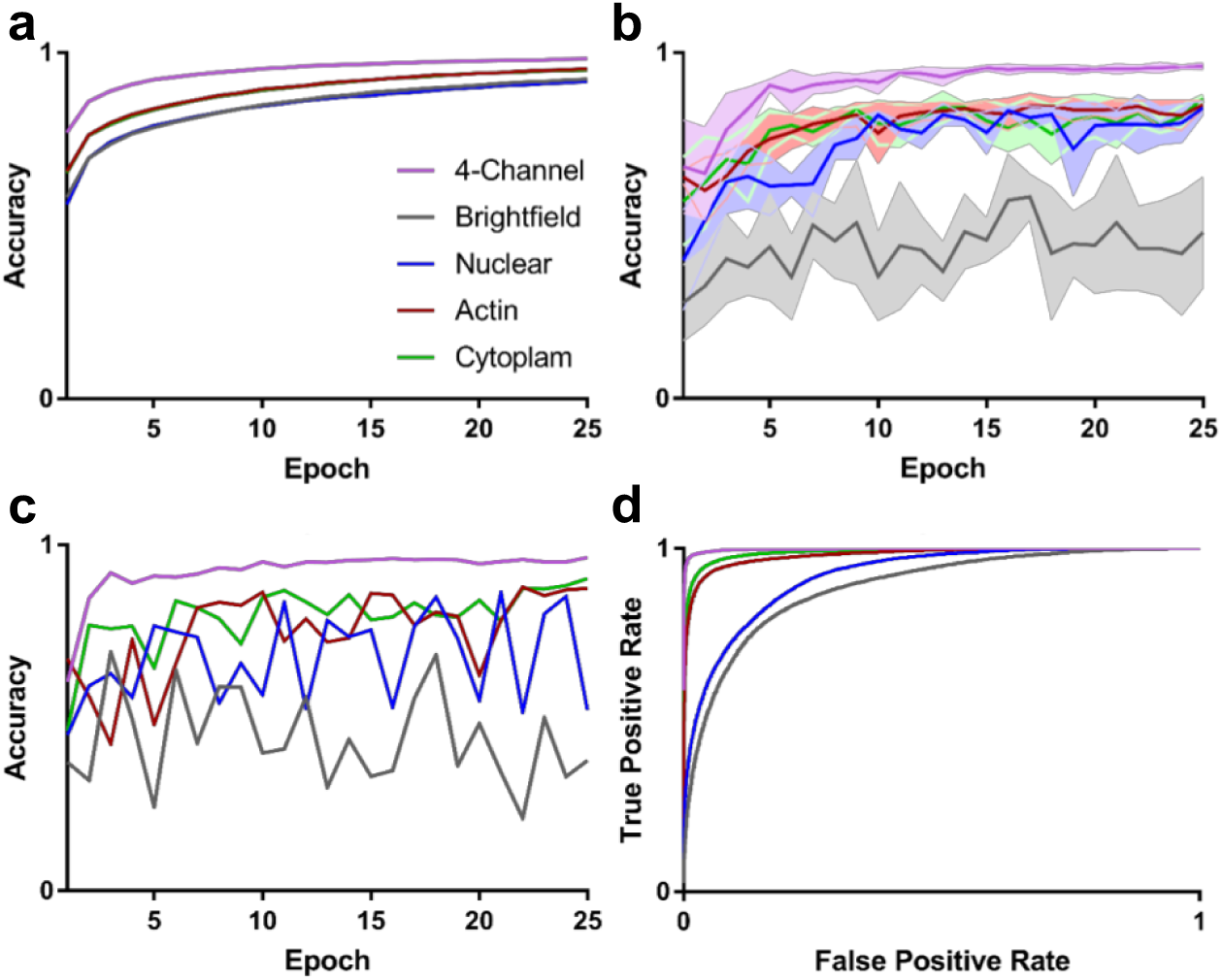
Training and Testing. **a,** Training accuracy results of the five models. Each model was trained over 25 epochs on a balanced dataset of 80,000 images. **b**, 5-fold exhaustive cross-validation results of the five models. The four-channel model reached a classification accuracy 0f 96% with a standard deviation of 0.8% after 25 epochs. **c,** Testing results of each of the five models, tested on 500 images of each class, sampled from a new dataset that was separately prepared and imaged. The four-channel model achieved classification accuracy of 96.3%. **d**, ROC curve displaying the macro-average trend across classes for each of the five models. The macro-average was computed from the results of each individual classes ROC curve, calculated using a one-vs-all approach.

### Cross-validation

We initially validated our results using exhaustive five-fold cross-validation. The training-set were randomly divided into five groups. The CNN was trained on four of the groups and tested on the fifth at the end of each epoch. The process was repeated four times, once for each combination. The five-fold cross-validation accuracy for the four-channel model plateaued after 25 epochs with an average accuracy of 96.0±0.8% (**Fig. 2b**). The three fluorescent channels achieved similar results with an average accuracy of 85.2±1.25%. The classification accuracy for the brightfield channel was significantly lower at 48.1±16.0%. The accuracy of the four-channel model combined with its low variability suggests a high level of confidence in the classification accuracy of the model.

### Testing

A key concern in assessing classification accuracy using five-fold cross-validation is the potential for batch effects where sampling artifacts are used by the model for classification.^25^ To address this concern, we separately prepared and imaged more cells from the same cell lines, and then classified them using our trained CNN’s. Each test set contained 500 randomly sample images from each class in order to obtain a balanced test set. Using this approach, we found the four-channel model achieved an accuracy of 96.3% (**Fig. 2c**), which is very similar to the cross-validation accuracy (**Fig. 2b**). The classification accuracy of the cytoplasm and cytoskeleton channels were 87.3% and 90.2%, which is also similar to cross-validation results. The nuclear channel had a much more volatile trend of testing accuracy compared to cross-validation suggesting there may have been some batch effects in the cross-validation results. The brightfield channel performed as expected with a final accuracy of 37.7%.

### Contributions from individual channels

To investigate the amount of information our method gains by incorporating fluorescent images we trained four networks using either the brightfield or one of the fluorescence channels and compared the classification accuracy with our four-channel model. The results show that the brightfield channel performed the worst out of all channels, implying that the channel had the least relevant information for classification (**Fig. 2b-d**). We believe this result stems from the brightfield channel only giving insight into the external morphology of cells, which is less distinctive because of disaggregation by trypsin, whereas the fluorescent channels provided insight into the internal structure of the cell. The nuclear channel was found to have the second lowest classification accuracy, which likely stemmed from the lower morphological diversity of nuclei. Finally, the cytoplasmic channel has the greatest single channel classification accuracy, which suggest the greatest amount of phenotypic information in this channel.

### ROC curve

To investigate how the model would perform in situations were the number of true positive or false positives is of great importance we computed a receiver operating characteristic (ROC) curve (**Fig. 2d**). The ROC curve was computed by analyzing the output probabilities of the model in a one-vs-all fashion, for each class, then computing the macro-average for each model. The resulting graph shows that the four-channel model had extremely high sensitivity and specificity. The ability to exchange sensitivity and specificity will be useful for applications, such as rare cell detection, that can tolerate some false positives to capture more true positives. In these results we can see that the cytoplasm and cytoskeleton models are also effective but would have to accept much higher fraction of false positives to achieve a similar yield of true positives.

### Confusion matrix

To summarize the classification accuracy results for each cell line, we plotted a confusion matrix together with the cumulative distribution for each cell line in each class to show the uniformity of the classification probabilities. The cumulative distribution function shows a histogram of the outputted class probability for all samples from each cell-line (**Fig. 3a**). The confusion matrix shows an average classification accuracy of 94.6% across all cell types, with HCT-116, HL-60, and PC3 cells reaching 99% accuracy (**Fig. 3b**). The lone exception was LNCaP, which had an accuracy of 84%, corresponding with the lowest number of training samples (2,555 samples).

**Figure 3.**
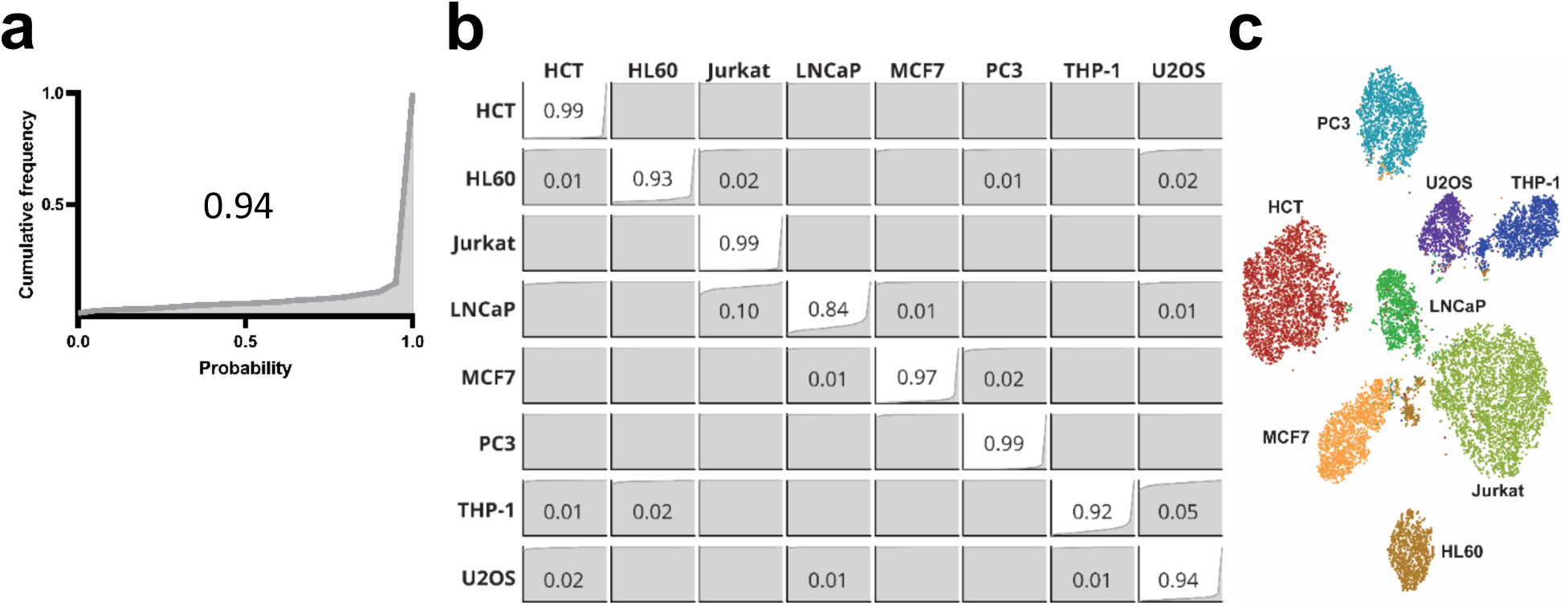
Classification accuracy and clustering. **a,** Example combined classification accuracy and cumulative probability distribution. The graph shows the cumulative probability frequency as a function classification probability for U2OS cells. **b,** Normalized confusion matrix constructed on the entire testing dataset. Diagonal values correspond to correct classifications, values <1% are not shown. Graphs in each position are the cumulative frequencies of classification probabilities for the corresponding classes. **c,** t-SNE graph computed from the 128-features extracted from the layer preceding the final classification layer for each image in the testing dataset.

### Clustering

We clustered the cell images via t-distributed stochastic neighbor embedding (t-SNE) plots. Using the output of the layer preceding the final output layer, we removed the final layer from the model and recorded the 128-feature output of the layer. We then used t-SNE with a perplexity of 64 and 2,000 iterations, to visualize this higher dimensional data in a two-dimensional graph (**Fig. 4**). The graph shows that the model is able to cluster the classes into individual clusters with significant separation. The quality of the clusters suggests that classical machine learning approaches could be used to separate the classes, using these features, with high confidence.

**Figure 4.**
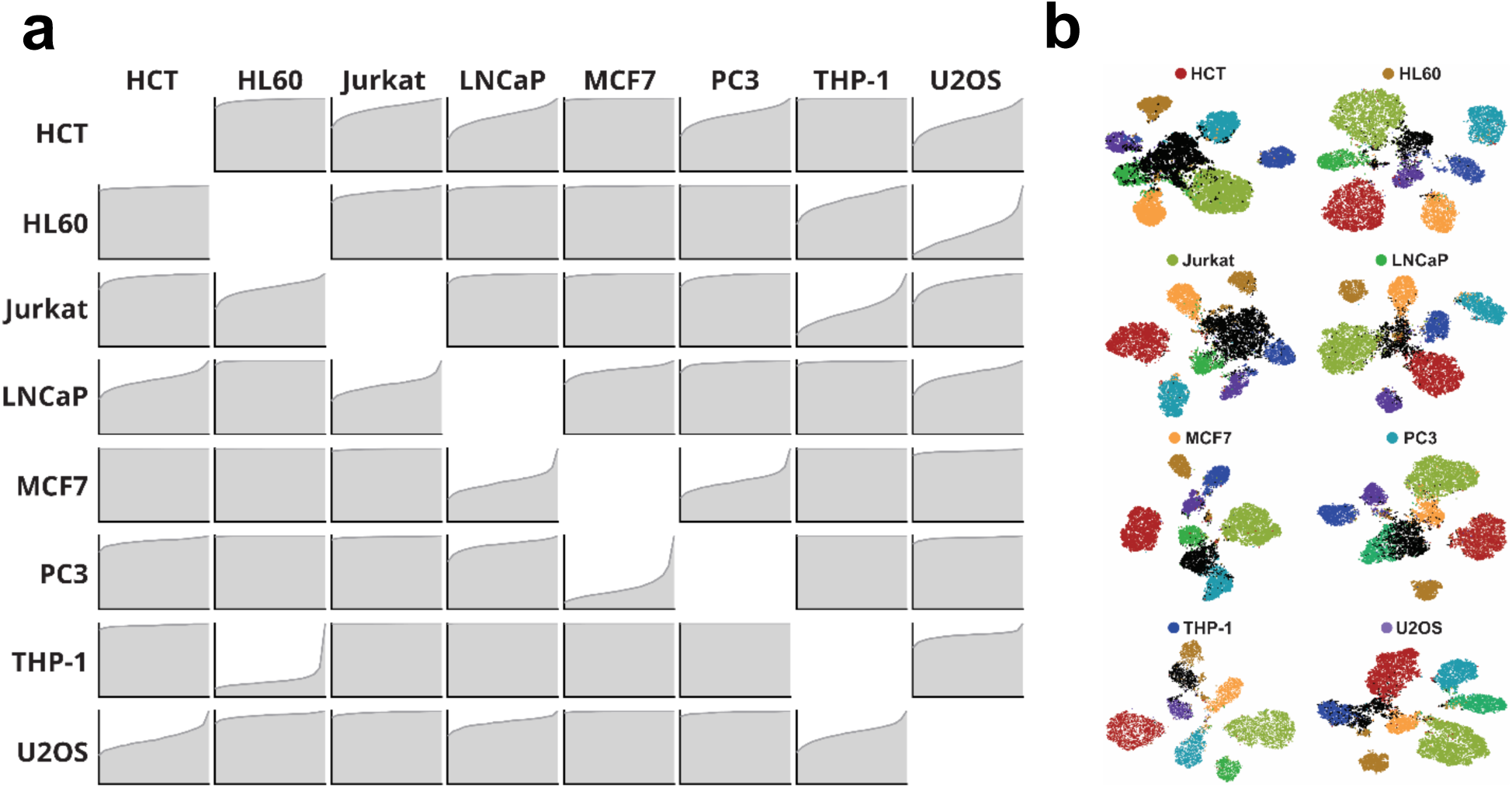
Classification and clustering of previously unseen cells. **a,** Classification probability distribution functions for each cell line when it is withheld from training. **b,** t-SNE plots of each cell line when it is withheld from training. The withheld class is shown in black, while the trained cell lines are labeled corresponding to **Fig. 3c**.

### Classifying and clustering previously unseen cells

To investigate how our CNN might respond to previous unseen cell types, we systematically designated a single cell line as previously “unseen cell line” and trained the CNN using only the seven other cell lines. We then classified the unseen cell line against these trained cells. This process was repeated by omitting each individual cell line and then classifying it against the remaining cells, allowing us to construct a matrix of cumulative classification probabilities (**Fig. 4a**).

These cumulative probability distributions show that previously unseen cell types are typically classified as one or two related cell lines. Interestingly, these queried cell lines were frequently classified as cell lines that are most closely related in cell lineage. For example, HL60 leukemia cell lines classified with the Jurkat and THP-1 leukemia cell lines. Similarly, MCF-7 classified as LNCaP and PC3 carcinomas. In some cases, the cell lines did not show lineage-specific classification. LNCaP classified with both Jurkat T-cell line as well as MCF-1 mammary adenocarcinoma line. However, this result may reflect the fact that LNCaP had the smallest training set and the lowest classification accuracy during the validation experiments. U2OS also classified with both leukemia and carcinoma cells but this likely reflects the fact that there were no other sarcoma cell lines for comparison.

Using the data generated by querying each cell line against the other seven cell lines, we visualized the clustering of each query cell line using t-SNE plots (**Fig. 4b**). In this visualization, the queried cell lines formed well-separated clusters from trained cell lines were generally located near cell lines of similar lineage. For example, clusters for MCF-7 cells were located near LNCaP and vice versa. Similarly, clusters for HL60, Jurkat, and THP-1 were located near each other.

## Discussion

In this study, we investigated whether deep-learning could phenotype single cells directly from microscopy images indistinguishable to the human eye. In contrast to traditional immunophenotyping where cell identities are determined using cell-specific antigen profiles, this study employed the distinct strategy of staining cells for common features, and then using deep-learning to distinguish cells from microscopy images. We examined eight cell lines that were dissociated by trypsin and stained with DAPI nuclear stain, Calcein-AM cytoplasmic stain, and SiR-actin cytoskeletal stain. Following unaided segmentation of microscopy images, we developed a CNN model that achieved a five-fold cross validation accuracy of 96.0±0.8%, as well as an average classification accuracy of 94.6% on a separate testing dataset. To determine whether the CNN model could be used to detect previous unseen cells, we queried each single cell type against a model generated by the other cell lines. While the efficacy of this approach varied between cell lines, it was remarkable that previously unseen cells were classified as cell lines in related differentiation lineages.

Analyzing microscopy images of trypsin-dissociated cells presented a significant challenge for image analysis because the enzymatic activity of trypsin protease causes cells to adopt a common spherical morphology. However, trypsin digestion also provide important practical advantages because disaggregated cells can be more evenly dispersed in a microscopy well-plate, which improves the robustness of the segmentation process. Previous studies to discriminate cells based on imaging required seeding cells on a surface, where segmentation is more complex and cell-surface interactions can influence cell morphology^26–28^. Disaggregated cell samples provide a simpler, more rapid, and more uniform imaging condition. Therefore, a key contribution of this work is the finding that images of disaggregated cells contain sufficient morphological information necessary for phenotyping.

A major technical hurdle in robust cell phenotyping using deep-learning arises from batch errors, which result from CNNs being trained on imaging and processing artifacts that do not reflect the biology of the cell phenotype^25^. In order to minimize batch errors, we generated the training set for each cell line from three separately prepared and imaged samples, to allow our model to capture variations in staining efficiency and illumination. We then ensured that our results were not biased by testing our CNN on a fourth separately prepared and imaged dataset. This further round testing was important as it avoided the potential biasing in our five-fold cross-validation as each fold was generated by randomly sampling the training set.

Together, this work demonstrates a deep-learning strategy that can classify cells based on common cellular phenotypes, rather than specific antigen markers. This finding is important because immunophenotyping is an expensive and error-prone process, despite significant efforts in standardizing staining, data collection and automation of analysis^5^. We observed that the morphological differences between cells may not need to be apparent to the naked eye in order to be sufficient for cell classification. Consequently, we have developed an approach for using disaggregated cells, similar to the preparation employed in standard laboratory flow cytometry, to establish rapid and robust image-based cell phenotyping of cells. With this capability, it is possible to imagine developing models for the 200+ known cell types in the human body in order to detect previously unknown phenotypes or detect phenotypic shifts that occur because of disease or treatment.

## Methods

### Sample Preparation

A total of 8 cancer cell lines were used in this study. Adherent cell lines were prostate derived cancer cell lines PC3 (ATCC CRL-1435) and LNCaP (ATCC CRL-1740), breast cancer MCF7 (ATCC HTB-22), colon cancer HCT 116 (ATCC CCL-247) and bone osteosarcoma U2OS (ATCC HTB-96). Suspension cell lines were leukemic T-cell lymphoblast Jurkat E6.1 (ATCC TIB-152), acute myeloblastic leukemic HL-60 (ATCC CCL-240) and acute monocytic leukemia THP-1 (ATCC TIB-202). All adherent cells, except LNCaP, were cultured in Dulbecco’s Modified Eagle Medium (DMEM) with 4.5g/L D-Glucose and L-Glutamine (Gibco), supplemented with 10% Fetal Bovine Serum (FBS, Gibco) and 1X Penicillin/Streptomycin (P/S, Gibco). All other cells were cultured in RPMI Medium 1640 with L-Glutamine, supplemented with 10% FBS and 1X P/S. All cells were incubated in T-75 flasks (Corning) at 37°C with 5% CO2. When needed, the adherent cell lines were released from the flasks with Trypsin-EDTA (0.25%, Gibco) and then washed twice in complete media twice, prior to resuspension for staining. Suspension cells were also washed twice in complete media. After resuspension, cells were stained with 5μg/mL Hoechst 33342 (H3570, Invitrogen), 50pM SiR-actin (CY-SC001, Cytoskeleton), and Live Green, 2μg/mL Calcein AM (C1430, Invitrogen), incubated at 37°C with 5% CO2 for 1 hour and then washed twice in PBS. Cells were resuspended in PBS and aliquoted at low density into Greiner Sensoplate 96-well glass bottom multiwell plates (M4187-16EA, Sigma-Aldrich).

### Microscopy

Microscopy imaging was performed using a Nikon Ti-2E inverted fluorescence microscope. Images were acquired using a Nikon CFI Plain Fluor 10X objective and a 14-bit Nikon DS-Qi2 CMOS camera. Images were captured using four channels: brightfield with phase contrast, DAPI (Nikon C-FLL LFOV, 392/23nm excitation, 447/60nm emission and 409nm dichroic mirror), mCherry (Nikon C-FLL LFOV, 562/40nm excitation, 641/75nm emission and 593 dichroic mirror) and EGFP (Nikon C-FLL LFOV, 466/40nm excitation, 525/50nm emission and 495nm dichroic mirror). Illumination for brightfield imaging was performed using the built in Ti-2E LED. Epifluorescence excitation was performed using a 130 W mercury lamp (Nikon C-HGFI). Gain, exposure and vertical offset were automatically determined using built-in NIS functions to avoid user biasing. Cells were imaged in 96-well glass plates (supplier). The concentration of cells in each well were diluted down to approximately 1000 cells to ensure adequate spacing between adjacent cells. An automated procedure was run on NIS using the Jobs function to take 16 images, on each of the 4 channels, inside of each well. The images were exported from NIS to standard TIFF format.

### Segmentation

The TIFF files were segmented using a custom python script. The script begins by extracting cell locations using a global Otto-threshold on the DAPI channel (nuclear stain) followed by object labeling using the SciPy library. Images from each location are then checked for usability. Specifically, a 75×75 pixel bounding box is defined around the centroid of each detected object. The 75-pixel size was selected because it was sufficiently large to fit single cells from the cell lines imaged using a 10X objective. The proposed image patches are then put through a number of rejection tests. First the nuclear channel is thresholder using an Otsu threshold on the patch and the number of nuclei present is quantified. If numerous nuclei are detected, or found on the edge of the bounding box, the patch is rejected. The next test involves checking cell viability and counting cell bodies in the cytoplasm channel. This is done by running an Otsu threshold on the cytoplasm channel and then ensuring a minimal count of pixels is present. If an adequate number of pixels is not present, the cell is assumed to be dead. Object detection is then done using the watershed algorithm; if more than one cell body is present, or if the cell body is touching the bounding box, it is rejected. The actin channel was checked only for contact with the bounding box as it was possible for actin filaments to be non-connected in numerous areas of the cell body. Images from acceptable locations were normalized, between 0 and 1, and added to a list of images in a numerical array. The numerical array of images was serialized and saved in pickle format, which has a lossless compression. This process was automatically repeated for each stack of TIFF images in the database.

### CNN model

A Convolutional Neural Network (CNN), shown in **Fig. 1c**, was designed in Python using the Keras library in TensorFlow. The network was designed to accept a 4-channel input of size 75×75 pixels. The model started with a 256-channel convolution layer with a kernel size of 7×7 with a stride of 1. The next layer is a 2×2 max-pooling layer with a stride of 1. Next is a 128-channel convolution layer with a kernel size of 5×5 and a stride of 1 followed by another 2×2 max-pooling layer. After this is two 64-channel convolution layers in series, each with a kernel size of 3×3 and a stride of 1. These layers are followed by a max-pooling layer of 2×2 and stride of 1 before being flattened for connecting to the fully-connected layers. Each convolution layer was followed by batch normalization and ReLU activation. Three 128-node, fully connected layers, each with ReLU activation batch normalization and 20% dropout, were used to learn on the features extracted using the earlier convolutional layers. The output of the network consisted of eight nodes, one for each class, with a soft-max error function used for back propagation. Data augmentation, in the form of random integer multiplications of 90-degree rotations, was used to up-sample the classes that had under 10,000 samples for training to ensure even classes and non-biasing during training.

### Training environment

The software was run on a single computer operating Windows 10 with an Intel i7-8700K running at 3.70 GHz. There was 64GB of DDR4 RAM running at 3200MHz. The graphics card was an 8GB GTX 1080. Training was done in Python 3.6.8 utilizing the TensorFlow 1.13.1 library.

### Training

Each iteration of the network was trained on 25 epochs, with Adam optimization and a learning rate of 0.001. The soft-max function was used as an error function for the backpropagation. Several different networks were trained to show the effects of varying different parameters on the inputs. First, the dataset was sub-sampled through cross-validation to investigate the variability of classification accuracy as shown in **Fig. 2b**. Second, the input layer was modified to accept each individual image channel to quantify the dependency of training on each channel as shown in all of **Fig. 2**.

### Cross-validation

To ensure the validation accuracy was accurate, five-fold exhaustive cross validation was used. The cross validation was done by splitting the training-set into five groups. The network was then trained five times on training data consisting of four of the five groups. The fifth group in each training session was used to determine a validation score. The five validation scores were averaged at each epoch to compute the reported validation accuracies.

### Confusion matrix

To identify the miss-classifications between labels a confusion matrix was constructed. All of the images in the testing dataset were used. A class label was predicted using the CNN and compared against the true class label. The diagonal positions correspond to true labels, while the off-diagonal positions show miss-classifications. The matrix was normalized by the number of images in each class to represent the probabilistic accuracy.

### t-SNE visualization

To visualize the relative clusters in our data a t-SNE plot was constructed on all of the images in the testing dataset, using the second-to-last fully connected layer of the network. The layer consisted of 128 nodes, giving a 128-feature vector for each validation image. This required removing the final layer from the network, after training, and classifying all the testing images.

### ROC curve

To validate the sensitivity of the model a ROC curve was calculated on all of the images in the testing dataset, using the probability outputs of each class. The ROC curve was calculated for each class using the SciKit library, which required a one-vs-all classification method in which all the other class are treated as a single group being compared to the current class of interest. As the classes preformed similarly, the macro-average ROC curve was calculated and displayed. The ROC curves were summed together and divided by the number of classes to find the average accuracy.

